# A transmissible γδ intraepithelial lymphocyte hyperproliferative phenotype is associated with the intestinal microbiota and confers protection against acute infection

**DOI:** 10.1101/2021.07.22.453366

**Authors:** Luo Jia, Guojun Wu, Sara Alonso, Cuiping Zhao, Alexander Lemenze, Yan Y. Lam, Liping Zhao, Karen L. Edelblum

## Abstract

Intraepithelial lymphocytes expressing the γδ T cell receptor (γδ IELs) serve as a first line of defense against luminal microbes. Although the presence of an intact microbiota is dispensable for γδ IEL development, several microbial factors contribute to the maintenance of this sentinel population. However, whether specific commensals influence population of the γδ IEL compartment under homeostatic conditions has yet to be determined. We identified a novel γδ IEL hyperproliferative phenotype that arises early in life and is characterized by expansion of multiple Vγ subsets. Horizontal transfer of this hyperproliferative phenotype to mice harboring a phenotypically normal γδ IEL compartment was prevented following antibiotic treatment, thus demonstrating that the microbiota is both necessary and sufficient for the observed increase in γδ IELs. Further, we identified a group of unique gut bacteria represented by 5 amplicon sequence variants (ASV) which are strongly associated with γδ IEL expansion. Using intravital microscopy, we find that hyperproliferative γδ IELs also exhibit increased migratory behavior leading to enhanced protection against bacterial infection. These findings reveal that transfer of a specific group of commensals can regulate γδ IEL homeostasis and immune surveillance, which may provide a novel means to reinforce the epithelial barrier.

## Introduction

Intraepithelial lymphocytes (IEL) are located within the intestinal epithelium and provide the first line of defense against luminal microorganisms^1^. Nearly half of murine IELs express the γδ T cell receptor (TCR), which exhibit a largely protective response to dampen acute inflammation^2, 3^ and promote mucosal barrier integrity^4, 5^. These protective functions have largely been attributed to IELs bearing the Vγ7 TCR, which comprise the majority of the γδ IEL population^6^. We have shown that γδ IELs limit microbial translocation by migrating into lateral intercellular space (LIS) between adjacent epithelial cells to provide surveillance of the barrier^7,8^. However, our understanding of the factors involved in regulating γδ IEL migratory behavior and activation remains limited.

γδ IELs are maintained in a partially-activated state to provide an immediate defense against invasive microbes, while limiting the potential for autoimmunity. Recently, activation of γδ IELs was shown to induce the production of interferons (IFN)^4^, which are potent immunomodulatory cytokines that are rapidly induced in response to viral or bacterial infection^9^. Type I IFN activates the IFNα/β receptor that is comprised of IFNAR1 and IFNAR2 (IFNAR). The presence of commensal bacteria at birth induces tonic type I IFN production by myeloid cells in the intestinal mucosa^10^, which contributes to the development of mucosal immunity^11, 12^. Although tonic IFNAR signaling is critical for the maintenance of lamina propria lymphocyte populations^13^, its effect on IELs, and γδ T cells in general, remains unclear.

Gnotobiotic, or germ-free (GF), mice exhibit an overall reduction in IELs, yet the number of γδ IELs remains largely intact^14, 15^. While these findings indicate that microbiota is not required for γδ IEL development, four-week-old GF mice exhibit a delay in population of the IEL compartment compared to SPF mice^16^. Further, administering antibiotics to wildtype (WT) mice immediately after birth significantly reduces γδ IEL number in the small intestine without affecting other peripheral lymphocytes^17^. Consistent with this, signaling through pattern recognition receptors and the aryl hydrocarbon receptor contribute to IEL homeostasis^17–19^. Whereas *Lactobacillus reuteri* promotes the development of CD4^+^ CD8αα^+^ IELs^20^, the extent to which specific commensal bacteria influence γδ IEL number and function has yet to be determined.

In this study, we have serendipitously discovered a novel γδ IEL hyperproliferative phenotype that arises early and persists throughout life. This γδ IEL expansion is driven by active proliferation of all Vγ subsets; however, the overall composition of this population skews toward Vγ7^−^ lymphocytes. We find that the microbiota is both necessary and sufficient to transfer the γδ IEL hyperproliferative phenotype to non-phenotypic mice, and further, we identified 5 amplicon sequence variants (ASV) that are closely associated with the expansion of γδ IELs. Interestingly, the hyperproliferative γδ IELs also exhibit enhanced migratory behavior at steady-state and confer protection against systemic *Salmonella* infection. These findings highlight the contribution of a specific group of commensals in the regulation of γδ IEL homeostasis and surveillance behavior, which may provide a novel means to reinforce the mucosal barrier in response to injury or infection.

## Results

### Commensal bacteria promote γδ IEL surveillance behavior

Previous studies have demonstrated that the microbiota is dispensable for population of the γδ IEL compartment; however, in the absence of commensal microbiota γδ IEL surveillance was reduced along the crypt-villus axis^21^. To determine the effect of the microbiota on the kinetics of γδ IEL migratory behavior within the epithelial compartment, GFP γδ T cell reporter mice (TcrdEGFP) were treated with broad-spectrum antibiotics and intravital microscopy was performed. We find that depletion of commensal bacteria results in reduced γδ IEL migratory speed, the frequency of migration into, and dwell time within the lateral intercellular space (LIS) (Fig. S1). These data support previous findings that commensal-derived signals contribute to γδ IEL motility patterns within the intestinal mucosa.

### Identification of a γδ IEL hyperproliferative phenotype that is accompanied by skewed Vγ subset composition

We and others have shown that epithelial-immune crosstalk is required to promote γδ IEL homeostasis and enable a rapid response to microbial pathogens. For example, epithelial MyD88 signaling promotes increased γδ IEL migration in response to bacterial infection^21^. Based on the known roles for tonic type I IFN in priming the host response to enteric infection^10–12^, we investigated the contribution of type I IFN signaling to γδ IEL homeostasis. To this end, we crossed IFNAR-deficient mice to those expressing the TcrdEGFP reporter. Morphometric analysis of the jejunum revealed a substantial increase in the number of GFP^+^ γδ IELs in adult IFNAR KO mice compared to TcrdEGFP (WT) (Fig. 1a,b). We find that this increase is due to enhanced γδ IEL proliferation as IFNAR KO mice exhibit a 50% increase in EdU^+^ γδ IELs relative to WT (Fig. 1c). Based on these findings, we next asked whether this enhanced proliferation could be attributed to a specific Vγ subset within the IEL compartment. Compared with WT, we were surprised to find that the relative proportion of γδ IELs skewed toward Vγ7^−^ subsets in IFNAR KO mice (Fig. 1d, S2a,b). Further analysis of each Vγ subset revealed that IFNAR KO mice exhibit a 22% and 66% increase in proliferation in Vγ7^+^ IEL and Vγ7^−^ IEL populations, respectively, compared to WT counterparts (Fig. 1e, S2c). Although the overall number is increased, the relative proportion of IEL subsets remained similar between WT and IFNAR KO mice (Fig. S2d). Interestingly, we found that both the frequency and the total number of γδ T cells from the spleen and mesenteric lymph nodes (MLN) were similar between the two genotypes (Fig. S2e,f).

**Figure 1.**
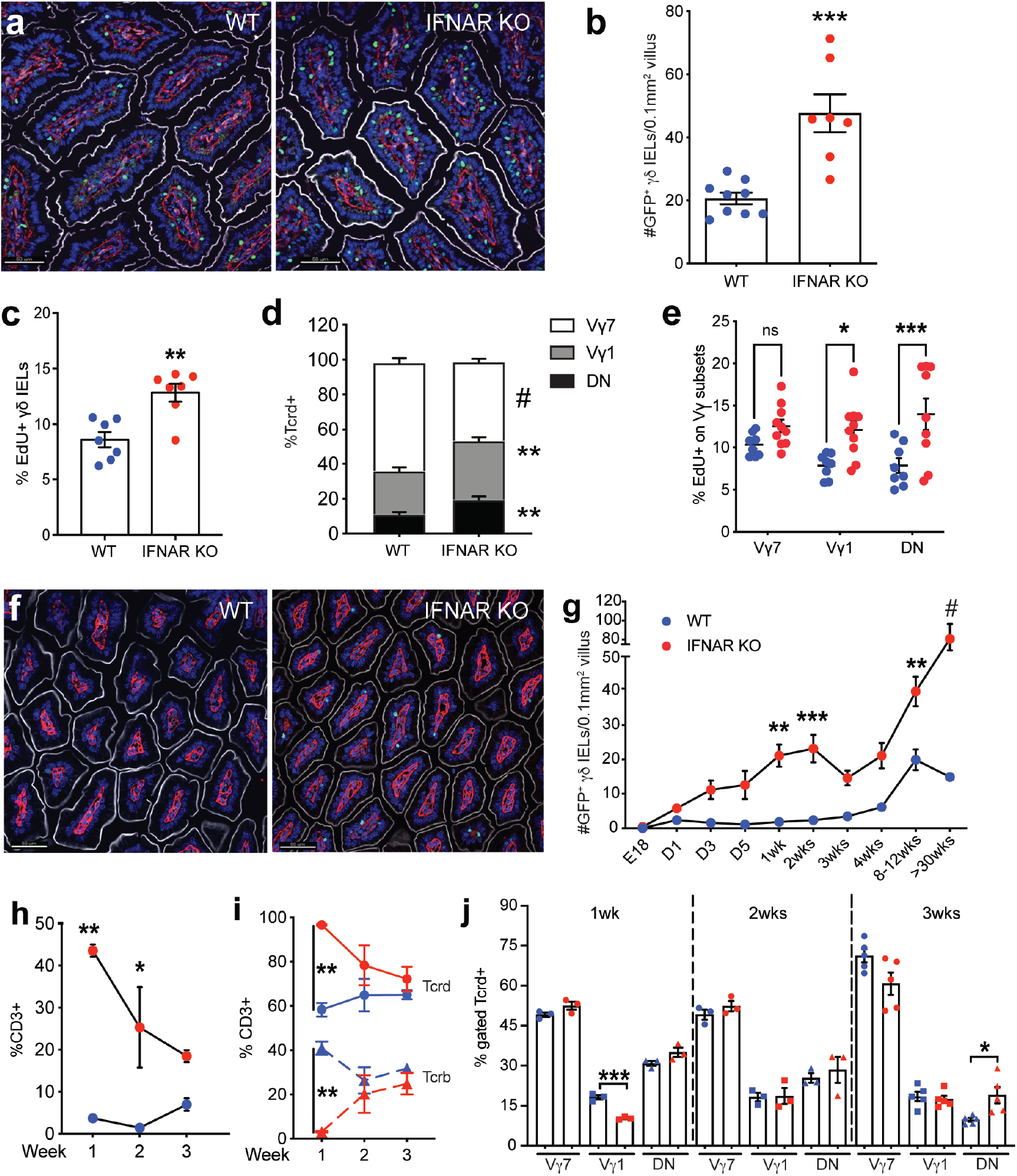
IFNAR-deficient mice exhibit a γδ, IEL hyperproliferative phenotype and skewed Vγ composition early in life. **a** lmmunofluorescent micrographs of jejunum from TcrdEGFP WT and IFNAR KO mice. Scale bar= 50μm. Nuclei, blue; laminin, red; γδ T cell, green; F-actin, white. **b** Quantification of jejuna GFP^+^ γδ T cells. **c** Percentage of EdU^+^ γδ IELs and **d** frequency of small intestinal Vγ IEL subsets in WT and IFNAR KO mice (gated on GFP). **e** Percentage of EdU^+^ cells (gated on individual Vγ subsets). **f** lmmunofluorescent micrographs of jejunum from 1-week old TcrdEGFP WT and IFNAR KO mice. Scale bar= 50μm. **g** Number of GFP^+^ γδ T cells in the small intestine of WT or IFNAR KO mice at various ages, n=4-10. Frequency of **h** CD3^+^ cells (gated on live cells), **i** Tcrd^+^ and Tcrb^+^ IEls (gated on CD3) and **j** Vγ subsets (gated on GFP) in WT and IFNAR KO postnatal mice, n=3-6. Data represent mean (±SEM) and are from at least two independent experiments. Each data point represents an individual mouse. Statistical analysis: **b,c,j:** unpaired t-test; **d,e,g,h:** two-way ANOVA with Sidak’s post hoc test; **i:** two-way AN OVA with Tukey’s post hoc test. *P<0.05, **P<0.01, *** P<0.001, # P<0.0001. ns, not significant.

To determine when this γδ IEL hyperproliferative phenotype is established, we performed morphometric analysis of intestinal tissue sections obtained from neonatal and weanling mice (Fig. 1g). As early as one week after birth, we observed a substantial increase in γδ IELs in IFNAR KO mice compared to WT (Fig. 1f,g). At one week of age, the emerging IEL compartment in IFNAR KO mice was heavily skewed toward TCRγδ^+^ IELs (Fig. 1h,i). However, the relative frequency among Vγ subsets was similar between the two genotypes at these early timepoints (Fig. 1j), suggesting that the differential expansion of Vγ subsets that we observed in IFNAR KO mice occurs post-weaning. These data demonstrate that the expansion of the γδ IEL compartment occurs early in life and continues throughout adulthood.

We next investigated the potential factors that could drive the expansion of and/or skewing of Vγ IEL subsets. Butyrophilin-like (Btnl) -1 and -6 expressed in the murine small intestinal epithelium jointly regulate the maturation and expansion of Vγ7^+^ IELs in the gut^22^. The expression of three major *Btnl* genes *(−1, −4, −6)* was similar in small intestine of neonatal and adult WT and IFNAR KO mice (Fig. S2g). IL-7 and IL-15 have known roles in the development and proliferation of intestinal γδ T cells^23, 24^, yet no change in expression was detected in the small intestine of adult mice (Fig. S2h,i). Although the ontogeny of γδ IELs is somewhat controversial with studies demonstrating that IELs develop both extrathymically and from thymic precursors^22^, we found no difference in DN2 and DN3 T cell precursors or γδ T cells in E18.5 fetal thymus (Fig. S2j,k). Taken together, these data indicate that the γδ IEL hyperproliferative phenotype develops after birth but cannot be attributed to altered butyrophilin or cytokine expression, or thymic γδ T cell development.

We next asked whether the increase in the γδ IEL compartment in IFNAR KO mice could be attributed to clonal expansion. Analysis of the γδ TCR repertoire revealed both WT and IFNAR KO γδ IELs are comprised of heterogeneous populations (Fig. S3). Consistent with our flow cytometric analysis (Fig. 1d), Vγ7, Vγ1, and Vγ4 subsets were the main TCRs expressed and Vγ7^−^ subpopulations were predominant in IFNAR KO mice (Fig. S3d). From these data, we conclude that under these highly proliferative γδ IELs maintain their characteristic heterogeneity.

### Horizontal transfer of the microbiota is necessary and sufficient to induce the γδ IEL hyperproliferative phenotype

The experiments described above were performed in a standard SPF barrier facility (S); however, a colony of IFNAR KO mice were also maintained in an enhanced barrier facility (E) in which murine norovirus and *Helicobacter* species are excluded. In this enhanced facility, TcrdEGFP (WT)-(E) and IFNAR KO-E mice were crossed resulting in the generation of WT, IFNAR^+/−^ and IFNAR-deficient littermates expressing the GFP γδ T cell reporter. To our surprise, morphometric analysis and flow cytometry revealed that IFNAR KO-E mice did not display the γδ IEL hyperproliferative phenotype; these mice exhibited a similar number of γδ IELs and proportion of Vγ subsets compared to WT-S and WT-E mice (Fig. 2a, S4). These findings provided the first evidence that loss of type I IFN signaling alone was not sufficient to induce the γδ IEL hyperproliferative phenotype.

**Figure 2.**
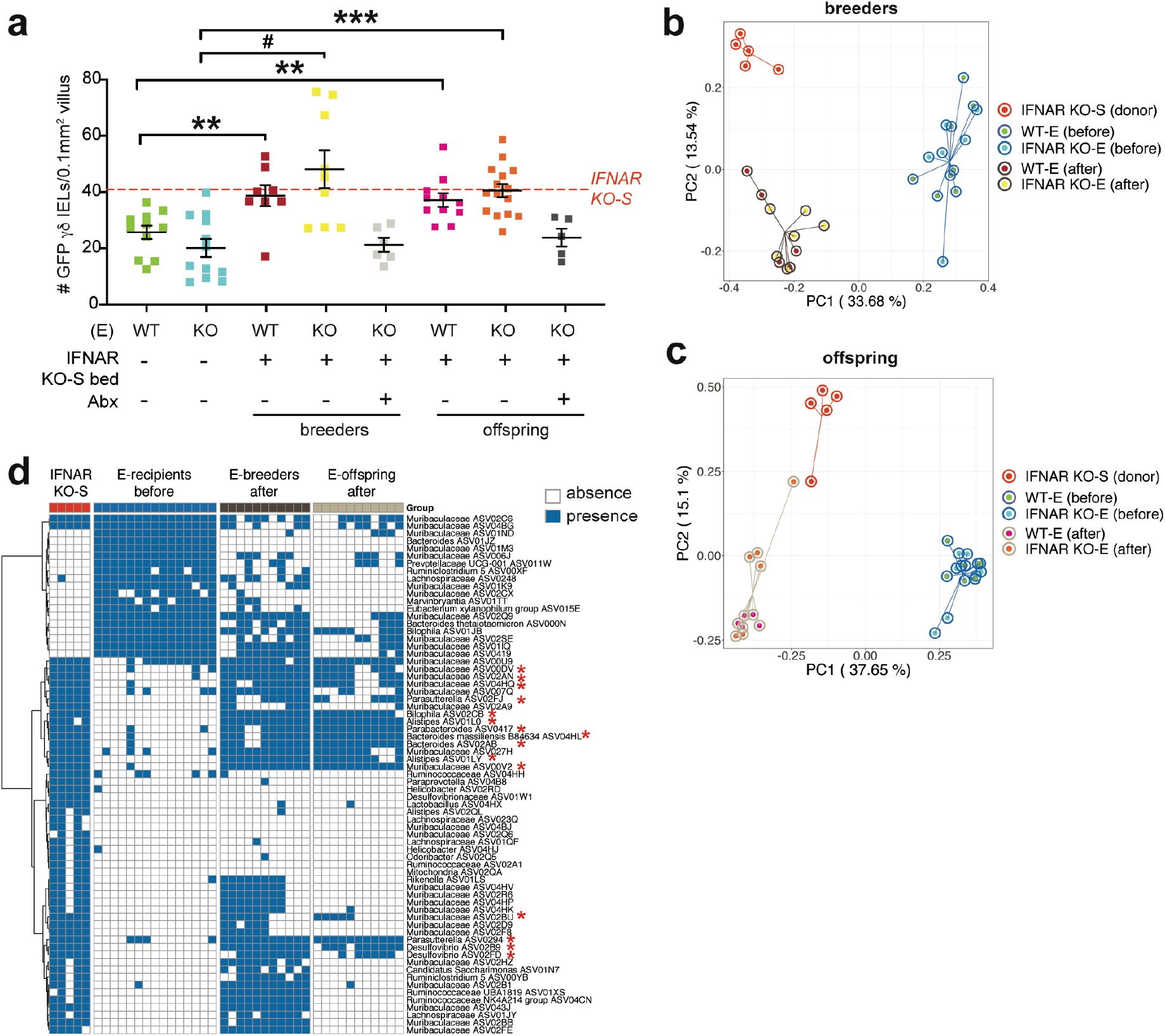
Horizontal transfer of the microbiota is necessary and sufficient to induce the γδ IEL hyperproliferative phenotype. **a** Morphometric analysis of the number of GFP^+^ γδ T cells in untreated WT-E, IFNAR KO-E mice; WT-E or IFNAR KO-E breeders and offspring following IFNAR KO-S bedding transfer in the presence or absence of antibiotic (Abx) treatment. Dashed line indicates the number of γδ IELs in IFNAR KO-S donor mice. n=5-15. Principal coordinates analysis of 16S rRNA sequencing of fecal microbiota from donor and recipient breeders **b** and offspring **c** n=4-6. **d** Binary heatmap of 69 ASVs shared in recipient mice (breeders and offspring) following horizontal transfer of microbiota. The red asterisk highlights the 15 ASVs which had significantly lower prevalence in pre-transfer breeders compared with donors, post-transfer breeders and post-transfer breeders’ offspring. Each data point represents an individual mouse. Data represent mean (±SEM). Statistical analysis: **a:** one-way ANOVA with Dunnett’s post hoc test **P<0.01, *** P<0.001, # P<0.0001; **d:** Fisher’s exact test *P<0.05

It is well-established that the microbiome can influence the immune phenotype of mice maintained in separate facilities^25^. Thus, we next investigated the contribution of the microbiota to the observed changes in the γδ IEL compartment. First, we asked whether this γδ IEL hyperproliferative phenotype could be transferred horizontally. Since mice are coprophagic, dirty bedding was transferred from cages of IFNAR KO-S mice into those housing breeding pairs of WT-E or IFNAR KO-E mice. After bedding transfer, the number of γδ IELs in WT-E or IFNAR KO-E breeders and their adult offspring resembled that of IFNAR KO-S mice (Fig. 2a). To determine whether the microbiota was required for the transfer of the γδ IEL hyperproliferative phenotype, broad-spectrum antibiotics were administered in parallel to the bedding transfer in a subset of cages. Antibiotic treatment was able to prevent the horizontal transfer of the γδ IEL phenotype to both WT-E and IFNAR KO-E breeders and their adult offspring (Fig. 2a). Collectively, these data demonstrate that an altered microbiota is both necessary and sufficient for horizontal transfer of the γδ IEL hyperproliferative phenotype.

To explore the changes to gut microbiota following horizontal transfer and identify the members associated with the γδ IEL hyperproliferative phenotype, we next performed 16S rRNA gene V4 sequencing on fecal samples collected from bedding transfer donors (IFNAR KO-S) and recipients (WT-E and IFNAR KO-E breeders and offspring). Prior to bedding transfer, the alpha diversity of WT-E and IFNAR KO-E breeders differed from that of IFNAR KO-S mice (Fig. S5a,b) which had higher richness. After bedding transfer, the alpha diversity of WT-E and IFNAR KO-E breeders was significantly higher than prior to transfer and more similar to that of IFNAR KO-S mice. Principal coordinates analysis (PCoA) based on Bray-Curtis distance showed a clear separation of the overall gut microbiota structure between donors and pre-transfer breeders along PC1 (Fig. 2b). Following bedding transfer, the gut microbiota of recipients became more similar to donors along PC1 though there was a separation along PC2. PERMANOVA test showed significant difference (P<0.001) between donors, pre-transfer and post-transfer breeders when compared in a pairwise fashion. Using redundancy analysis (RDA) with the group information as constraining variable, 101 ASVs were identified as differential variables between donors, pre- and post-transfer breeders (Fig. S6a, S7a). Next, we performed the same analysis between donors, pre-transfer breeders and the offspring of post-transfer breeders. The alpha diversity of the gut microbiota in the offspring was significantly lower than that of the other two groups (Fig. S5c,d) and their overall gut microbiota structure was more similar to donors compared to their parents prior to transfer (Fig. 2c). PERMANOVA test showed a significant difference between the three groups (P<0.001) and 108 ASVs were identified as differential variables using RDA (Fig. S6b, S7b). After combining the results from the two analyses, 69 ASVs were commonly identified by RDA (Fig 2d, S6c), from which 15 ASVs showed significantly lower prevalence in pre-transfer breeders as compared with donors, post-transfer breeders and their offspring. There was no marked difference between the latter three groups. Taken together, these analyses identified 15 ASVs that are associated with the horizontal transfer of the γδ IEL hyperproliferative phenotype.

### The γδ IEL hyperproliferative phenotype can be transmitted vertically to WT offspring

Our findings indicate that the microbiota, and not loss of IFNAR signaling, may be the primary factor leading to the expansion of the γδ IEL compartment. When initially generating the IFNAR KO-S line, we crossed the parental TcrdEGFP (WT-S) to IFNAR KO-S mice and subsequently housed the two strains separately. Housing mice of different genotypes separately can induce a confounding variable when evaluating the effect of the microbiome on a given phenotype. Thus, to control for the maternal effect of the microbiome, we crossed separately housed WT-S dams with IFNAR KO-S sires to generate F2 littermates (Fig. 3a). We observed that adult F2 WT mice exhibited a 2-fold increase in the number of γδ IELs, similar to parental IFNAR KO-S mice (Fig. 3b,c). Further, F2 WT γδ IELs exhibit enhanced proliferation compared to WT-S (Fig. 3d). Vγ1^+^ cells were increased and Vγ7^+^ cells were largely decreased in all three genotypes of F2 littermates relative to WT-S mice (Fig. 3e). The reciprocal cross was performed (Fig. S8a) and the resultant F2 WT mice showed a similar expansion of the γδ IEL compartment (Fig. S8b-d), indicating that the genotype of the dam was not a contributing factor. Together, these data demonstrate that the observed phenotype is vertically transmissible and confirms our earlier observation that γδ IEL hyperproliferation occurs independently of IFNAR signaling.

**Figure 3.**
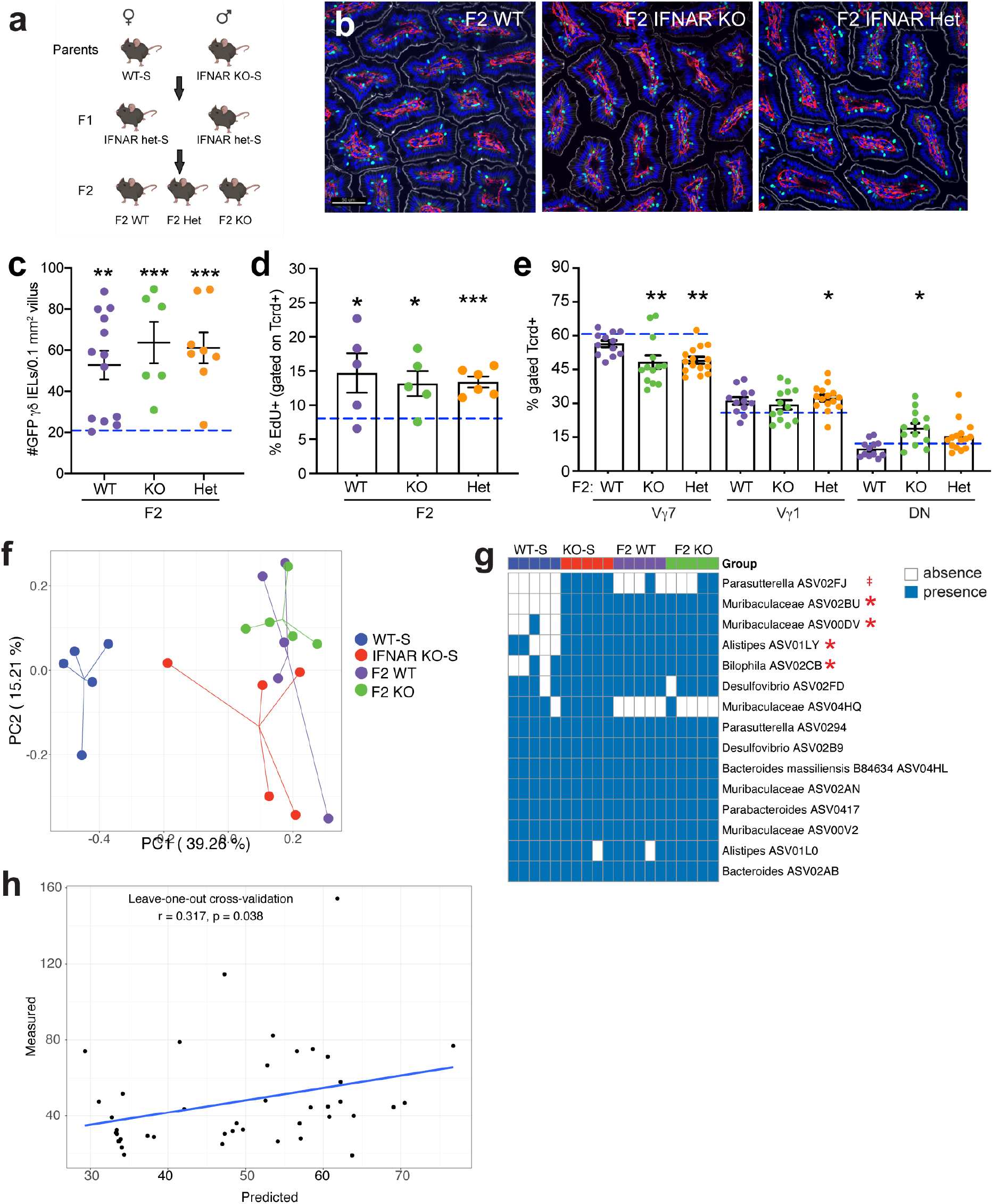
The γδ IEL hyperproliferative phenotype is transmitted vertically independently of genotype. **a** Breeding strategy used to generate F2 WT and IFNAR KO littermates. **b** Representative images and **c** morphometric analysis of GFP+ γδ T cells in the jejunum of adult F2 littermates. Scale bar= 50μm. n=6-13 mice in three independent experiments. **d** Percentage of EdU+ γδ IEls and **e** proportion of Vδ IEL subsets in adult F2 littermates. Dashed lines indicate separately housed WT-S values. **f** Principal coordinates analysis of 16S rRNA sequencing of fecal microbiota from WT-S, IFNAR KO-S and F2 WT and IFNAR KO littermates. **g** Occurrence heatmap of the 15 ASVs identified in the horizontal transfer dataset. Blue and white show the presence or absence of ASVs, respectively. **h** Associations between the 5 ASVs (highlighted in **g)** and the γδ IEL hyperproliferative phenotype. Data represent mean (±SEM) from two independent experiments. Each data point represents an individual mouse. Statistical analysis: **c,d:** unpaired t test compared to separately housed WT-S values; **e:** one-way ANOVA with Dunnett’s post hoc test; **g:** Fisher’s exact test was performed to compared prevalence of ASVs between WT-Sand the combination of the other 3 groups. **h:** Random Forest model with leave-one-out cross-validation was applied to use the ASVs abundance to regress the γδ IEL cell number. Pearson correlation was used to compare the predicted and measured values. ‡P<0.1 *P<0.05, **P<0.01, *** P<0.001.

Microbiome analysis of fecal samples collected from WT-S, IFNAR KO-S, F2 WT and F2 IFNAR KO mice revealed different alpha diversity of gut microbiota among the 4 groups (Fig. S9). Specifically, WT-S had significantly lower richness compared to the mice exhibiting the γδ IEL hyperproliferative phenotype, and the alpha diversity of the gut microbiota in F2 littermates was similar to IFNAR KO-S mice. PCoA plot based on Bray-Curtis distance showed a clear separation between WT-S and the phenotypic groups along PC1 (Fig. 3f) (PERMANOVA test, P < 0.001 between WT-S and the other 3 groups). The F2 littermates of both genotypes had a similar gut microbiota composition (PERMANOVA test, P = 0.86) close to IFNAR KO-S along PC1, but still showed a significant difference along PC2 (PERMANOVA test, F2 IFNAR KO vs IFNAR KO-S: P = 0.02; F2 WT vs IFNAR KO-S: P = 0.14). We next assessed the prevalence of the 15 bacterial ASVs identified in the horizontal transfer datasets (Fig. 2d) in this experimental group. Among the initial 15 ASVs, we found 5 (one from *Parasutterella*, two from *Muribaculaceae*, one from *Alistipes* and one from *Bilophila*) with significantly higher prevalence in the phenotypic mice compared to WT-S (Fig. 3g). We further explored the associations between these 5 ASVs and the number of γδ IELs using Random Forest regression with leave-one-out cross-validation. The predicted values showed a significantly positive correlation with the measured values indicating that these commensal bacteria likely contribute to the γδ IEL hyperproliferative phenotype (Fig. 3h).

### Hyperproliferative γδ IELs exhibit enhanced surveillance behavior to confer protection against bacterial infection

We have previously reported that γδ IELs provide continuous surveillance of the intestinal epithelium to limit acute pathogen invasion^8, 26^. Thus, to determine whether the expansion and/or alteration of γδ IEL subsets affects their overall surveillance behavior, we performed intravital microscopy of WT-S and F2 WT small intestinal mucosa under homeostatic conditions. Image analysis of time lapse videos revealed that γδ IELs in WT mice with the hyperproliferative phenotype migrated into the LIS more frequently than those in WT-S mice (Fig. 4a,b, Video S1). This increase in flossing behavior was accompanied by increased average track speed and reduced dwell time in the epithelium of F2 WT mice compared to WT-S. (Fig. 4c,d). As expected, the frequency with which an enterocyte was contacted by a γδ IEL was increased due to the expansion of γδ IELs within the epithelial compartment (Fig 4e).

**Figure 4.**
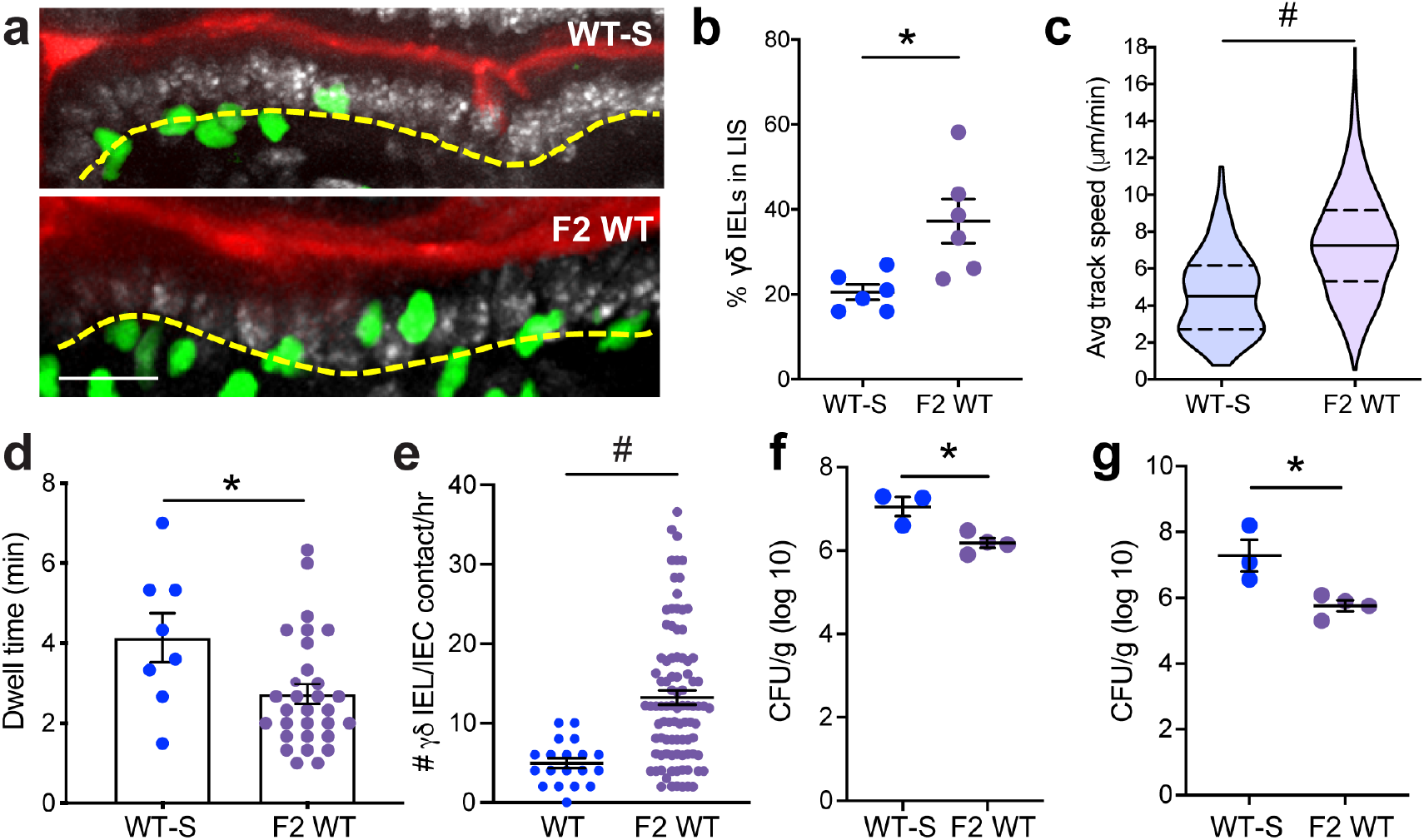
γδ IEL surveillance behavior is enhanced in WT mice exhibiting the hyperproliferative phenotype and confers protection against enteric infection. **a** Maximum projection images taken from intravital microscopy of γδ IEls (green) within the jejunal epithelium of WT-S and F2 WT mice. Scale bar= 30μm. Nuclei, white; lumen, red. Yellow dashed line approximates the basement membrane. **b** Frequency of γδ IEls in the lateral intercellular space (LIS), **c** average track speed, **d** dwell time and **e** γδ IEL/IEC contacts/hr. WT-S and F2 WT mice were infected orally with *Salmonella* Typhimurium. CFU were measured at 6 dpi in **f** spleen and **g** liver homogenates. Each data point represents an individual mouse, except **c,d,e** in which each data point represents an individual track, γδ IEL or IEC, respectively. Data represent mean (±SEM). Statistical analysis: unpaired t-test. *P<0.05, # P<0.0001.

We have previously demonstrated that migration of γδ IELs into the LIS is critical to limit *Salmonella* Typhimurium invasion and subsequent spread to peripheral sites^8^. To address whether the expansion and increased motility of γδ IELs contributes to the host response to infection, WT-S and F2 WT mice were infected orally with *Salmonella*. Six days post-infection, F2 WT mice showed reduced bacterial load in the spleen and liver compared to WT-S mice (Fig. 4f,g) demonstrating that the transfer of this hypermotile and hyperproliferative γδ IEL phenotype to WT mice results in enhanced γδ IEL surveillance capacity to effectively reduce the severity of systemic salmonellosis.

## Discussion

In this study, we serendipitously identified a novel group of commensals that correlate with the expansion of the γδ IEL compartment. This increase in proliferation is polyclonal and not restricted to one Vγ subset, although Vγ7^−^ IELs expand to a greater extent than Vγ7^+^ IELs. While we initially attributed this finding to a loss of tonic IFNAR signaling, differences in the IEL compartment between mice housed in different facilities pointed to an environmental factor as the potential cause. Through horizontal and vertical transfer experiments, we determined that the microbiota was required for this phenotype, and further identified 5 ASVs that correlate with the expansion of γδ IELs. In addition to increased proliferation, we found that these lymphocytes exhibit enhanced surveillance behavior within the epithelium. Consistent with our previous reports^8, 26^, we showed that transfer of the γδ IEL hyperproliferative phenotype to WT mice results in protection against systemic salmonellosis.

Microbial colonization early in life plays a key role in the development of mucosal immunity. Unlike MAIT cells, antigen-specific effector and regulatory T cells that develop during early life in response to commensal exposure^27–30^, the microbiota is dispensable for γδ IEL development^14, 15^. However, we find horizontal transfer of microbiota can induce the expansion of γδ IELs in both neonatal and adult mice. We now describe a group of 5 ASVs associated with enhanced γδ IEL proliferation including two from the family *Muribaculaceae*, and one from each of the following genera, *Parasutterella*, *Alistipes* and *Bilophila.* Notably, these ASVs are all anaerobic Gram-negative bacteria, which in combination with the adhesive and mucin-degrading properties of *Bilophila*^31^, may permit greater access to the epithelial surface and subsequently activate pattern recognition receptors. In support of this, TLR/MyD88 or NOD2 signaling has been shown to contribute to the maintenance of CD8αα^+^ IELs via IL-15 production^17, 32, 33^. However, we did not detect a difference in *Il15* expression in the mucosa of these mice (Fig. S2). Thus, the precise mechanism by which these commensals promote γδ IEL proliferation remains unclear.

Several factors contribute to the selection and maintenance of γδ T cell subsets at barrier surfaces. Interestingly, we did not observe altered Vγ populations in the small intestine in neonatal or weanling mice (Fig. 1j), indicating that the bias towards Vγ7^−^ IELs likely occurs post-weaning. The local microbiota drives the proliferation of Vγ6^+^ T cells in the oral mucosa, lung and reproductive tract^34–36^; therefore, we speculate that the continued expansion and stabilization of the local microbial community may induce the preferential proliferation of Vγ7^−^ IELs. While the reciprocal interactions between γδ17 cells and the microbiota have begun to be elucidated^34–36^, the how commensals influence γδIFN populations remains poorly understood.

To date, the only commensal that has been identified to be associated with the development of a specific IEL subpopulation is *L. reuteri^20^*. However, five members of the *Muribaculaceae* family were also associated with the development of CD4^+^ CD8αα^+^ IELs. Since members of *Muribaculaceae* are difficult to culture, their specific role in IEL development has not been directly tested. We find that two *Muribaculaceae* family members are associated with the γδ IEL hyperproliferative phenotype as is *Bilophila.* One species of this genera, *Bilophila wadsworthia*, has been shown to adhere to human intestinal epithelial cells and exhibits genotoxic and pro-inflammatory properties^31^. Of note, we did not observe clinical disease or distortion of the mucosal architecture in mice displaying the γδ IEL hyperproliferative phenotype. It was recently reported that the sulfide production by taurine-utilizing bacteria such as *Bilophila wadsworthia* can inhibit pathogen aerobic respiration^37^. Therefore, an increase in sulfide-producing commensals may not only induce colonization resistance, but our data now suggest a potential role for these bacteria in modulating IEL-mediated surveillance. Interestingly, laboratory mice that have been “rewilded” or housed in a natural, outdoor environment also exhibit an increase in *Bilophila*^38^. Moreover, microbiome analysis of wild mice revealed an increase in order *Desulfovibrionales* and genus *Alistipes* in the gut that was accompanied by a concomitant increase in lamina propria CD8^+^ T cells. These wild mice showed improved clearance of infection^39^. While the murine IEL compartment has yet to be investigated under these natural housing conditions, taken together, these studies raise the possibility that an increase in these commensals may improve the host mucosal immune response to invasive pathogens.

Although our data demonstrate that the γδ IEL hyperproliferative phenotype occurs independently of IFNAR signaling, we cannot rule out the possibility that loss of type I IFN signaling contributed to the expansion of specific ASVs. The results from our microbiome analyses are consistent with previous reports indicating that loss of global IFNAR signaling (IFNAR KO-E mice) does not induce significant changes to the fecal microbial community^40^. Interestingly, we find that the γδ IEL hyperproliferative phenotype correlates with increased *Alistipes spp,* which were also expanded in IEC-specific IFNAR KO mice^41^.

We find that the microbiota is both necessary and sufficient to enhance γδ IEL proliferation; however, we cannot exclude the effect of the microbiota on commensal viruses, which also contribute to IEL maintenance^18^. Murine norovirus (MNV) is present in our standard (S) but not enhanced (E) barrier facility. Whereas MNV can induce IFNAR signaling in the absence of an intact microbiota to maintain in lamina propria lymphocyte populations^13^, the role of MNV in the regulation of γδ IELs is unknown. Thus, the extent to which type I IFN signaling modulates the IEL compartment and the potential contribution of commensal viruses to this phenotype would be of interest for future investigation.

Dysregulation or aberrant expansion of the IEL compartment is associated with disease states such as celiac disease or inflammatory bowel disease^42^; however, we do not observe overt intestinal pathology in phenotypic mice. Although further study is required to better understand how this microbiota affects other aspects of mucosal immunity, our findings open new avenues to explore how the microbiota or microbial-derived products can be manipulated to modulate γδ IEL proliferation and migratory behavior as a means to reinforce the epithelial barrier in the context of gastrointestinal infection.

## Methods

### Animals

All mice were maintained on a C57BL/6 background and unless otherwise noted, mice of both sexes were analyzed between 8-12 weeks of age. Mice were housed under SPF barrier conditions, with colonies maintained in a standard barrier (S) or enhanced barrier facility (E) which is *Helicobacter*- and murine norovirus-free. Mice were fed an autoclaved commercial rodent 5010 diet and tap water. All mice were kept in the room with standard 12 hours light-dark cycle and humidity and temperature were monitored. TcrdEGFP mice^43^ were crossed to IFNAR1 KO mice (provided by Sergei Kotenko and Joan Durbin, Rutgers NJMS). F2 littermates were generated in the SBF by crossing separately housed WT-S and IFNAR KO-S mice. All studies were conducted in an Association of the Assessment and Accreditation of Laboratory Animal Care–accredited facility using protocols approved by Rutgers New Jersey Medical School Comparative Medicine Resources.

### Intravital microscopy

Time lapse intravital microscopy of the jejunal mucosa was performed as previously described^44^. Image analysis was performed using Imaris (v.9.7; Bitplane), in which surface-to-surface distance was calculated between GFP^+^ γδ T cells and the lumen. γδ T cells within 10-12 μm from the lumen were considered within the LIS. γδ IEL track speed was calculated by an autoregressive tracking algorithm confirmed by manual verification of individual tracks. Dwell time and the frequency of γδ IEL/epithelial interactions were quantified manually.

### *In vivo* treatments

Vancomycin (400 μg/mL, Hospira) meropenem (200 μg/mL, Bluepoint Laboratories) were administered in the drinking water for two weeks. For bedding transfer experiments, WT-E and IFNAR KO-E breeding pairs that were transferred into the SBF (S) facility were housed with dirty bedding obtained from IFNAR KO-S cages, which was replenished biweekly. Some breeding cages were maintained on Abx while exposed to dirty bedding (Fig. 2). All offspring were weaned into clean cages with tap water. Mice were injected intraperitoneally with 0.5 mg 5-Ethynyl-2’-deoxyuridine (EdU, TCI America) daily for five days. Mice were gavaged with 200 mg/ml streptomycin (Sigma-Aldrich) 24 h prior to gavage with 10^8^ colony-forming units (CFU) of DsRed-labeled *Salmonella Typhimurium* (strain SL3201; Andrew Neish, Emory). Mice were euthanized six days post-infection after which spleen and liver homogenates were plated and colonies counted.

### Immunofluorescence and image analysis

Neonatal and post-neonatal (0-3 weeks old) intestine was fixed in 10% formalin and embedded in paraffin, whereas post-weanling and adult intestine were fixed and embedded as described previously^26^. Briefly, tissue was fixed in 1% paraformaldehyde, washed with 50 mM NH4Cl, cryoprotected in 30% sucrose (wt/vol) and embedded in Optimal Cutting Temperature (OCT, Tissue-Tek) medium. Immunostaining of frozen or FFPE sections (5-7 μm) was performed using the rabbit anti-laminin (Sigma-Aldrich) or biotin-labeled anti-GFP (Abcam), followed by Alexa Fluor 594 goat anti-rabbit IgG (H+L), Alexa Fluor 647 phalloidin, Alexa Fluor 647 Streptavidin (Invitrogen) and/or Hoechst 33342 dye (Invitrogen). Slides were mounted with ProLong Glass (Invitrogen) and images were acquired on an inverted DMi8 microscope (Leica) equipped with a CSU-W1 spinning disk, ZYLA SL150 sCMOS camera (Andor), PL APO 40x/0.85 dry objectives, and iQ3 acquisition software (Andor). The number of GFP^+^ cells per 0.1 mm^2^ villus was quantified by an observer blinded to the condition.

### IEL isolation and flow cytometric analysis

Small intestinal IELs were isolated as previously described^7^ and GFP^+^ γδ IELs were sorted to 98% purity using a BD FACSAria II. IELs were stained with viability dye (eFluor 450 or eFluor 780), anti-CD3 (2C11), anti-CD8α (53-6.7), anti-CD8β (H35-17.2), anti-CD4 (GK1.5), anti-TCRβ (H57-597), anti-TCRγδ (GL3) (eBioscience), anti-Vγ7 (clone GL1.7; Rebecca O’Brien, National Jewish Health, Denver, CO), and anti-Vγ1 (clone 2.11, BioLegend) or a Click-iT Plus EdU Alexa Fluor 647 Flow Cytometry Assay Kit (Invitrogen). Flow cytometry was performed on an LSR Fortessa (BD Biosciences) in the New Jersey Medical School Flow Cytometry and Immunology Core Laboratory. Data were analyzed by FlowJo (v.10.4.0; Tree Star).

### TCR repertoire data analysis

RNA was isolated from 3×10^5^ γδ IELs, extracted using TRIzol (Invitrogen) and purified by RNeasy Mini Kit (Qiagen). Library construction and sequencing were performed by iRepertoire (Huntsville, AL, USA). The usage of V, D and J gene and complementarity-determining region 3 (CDR3) sequences were determined, and tree maps were generated using iRweb tools (iRepertoire). Datasets were processed using the MiXCR software package (v3.0.13) to further correct for PCR and sequencing errors. Diversity metrics, clonotype overlap and gene usage were plotted in R, by VDJTools (v1.2.1) using standard packages.

### qPCR assays

RNA was extracted from homogenized tissue as described above. Total RNA was reverse transcribed using an iScript cDNA Synthesis Kit (Bio-Rad) and qPCR was performed using SYBR Green (Fisher Scientific) using QuantStudio 6 platform (ThermoFisher Scientific). Ct values were normalized to the Ct values for *Ecad (Btln1,4,6,Il15)* and *Gapdh (Il7),* respectively.

### Gut microbiome analysis

Genomic DNA was extracted using protocol Q^45^ with minor modification. Hypervariable region V4 of the 16S rRNA gene was amplified using the 515F and 806R primers modified by Parada et al.^46^ and Apprill et al.^47^, respectively and sequenced using the Ion GeneStudio S5 (ThermoFisher Scientific). Primers were trimmed from the raw reads using cutadapt via QIIME 2. Amplicon sequence variants (ASVs) were obtained by denoising using the dada2 denoise-single command in QIIME 2 with parameters --p-trim-left 0 --p-trunc-len 215. Spurious ASVs were further removed by abundance filtering^48^. A phylogenetic tree of ASVs was built using the QIIME 2 commands alignment mafft, alignment mask, phylogeny fastree and phylogeny midpoint-root.

Taxonomy assignment was performed by the q2-feature-classifier plugin in QIIME 2 based on the silva database (release 132)^49^. The data were then rarified to 38,000 reads/sample for subsequent analyses. Alpha diversity indices (Shannon Index, observed ASVs, Faith’s phylogenetic diversity and evenness) and Bray-Curtis distance were used to evaluate the overall microbiota structure. Principal coordinates analysis (PCoA) was performed by the R packages “ape”. Redundancy analysis (RDA) which describes the relationships between the grouping and ASVs in microbial communities, was performed with R package “vegan” based on Hellinger transformed abundance of the ASVs. ASVs, which had more than 50% of the variation can be explained by the grouping, were selected. Random Forest analysis was performed and cross-validated using the R “randomForest” and “caret” package to test for correlations between ASVs and host phenotypes.

### Statistical analyses

Data were shown as the mean ± SEM. Statistical analysis was conducted in GraphPad Prism8. The significance between two independent samples was determined by unpaired t-test or for multiple independent variables one-way or two-way ANOVA was performed.

### Data and code availability

All raw 16S rRNA sequencing data is accessible in NCBISRA with accession number: PRJNA744534.The accession number of γδ TCR sequencing data is PRJNA744491.

## Acknowledgments

The authors would like to thank Sergei Kotenko and Joan Durbin for providing the IFNAR KO mice. Cell sorting was performed at the NJMS Flow Cytometry and Immunology Core Laboratory and supported by National Institute for Research Resources Grant S10RR027022. This work was supported by Busch Biomedical Research Grant, New Jersey Commission on Cancer Research Bridge Grant (DCHS19CRF009), National Institute of Health Grant DK119349 (K.L.E.).

## Author Contributions

L.J. designed and performed experiments and wrote the manuscript. G.W. analyzed data and wrote the manuscript. L.J. and G.W. contributed equally to the work. S.A. performed experiments and analyzed the data. A.L analyzed the data and C.Z. and Y.L. performed experiments. L.Z. contributed to experimental design, supervised data analysis and revised the manuscript. K.L.E. conceived the study, performed experiments, supervised the research and wrote the manuscript. All authors approved the final manuscript.

## Supplementary Video 1

Intravital microscopy of γδ T cells (green), luminal Alexa Fluor 633 (red) and nuclei (white) in jejunum of TcrdEGFP (WT) or F2 WT mice. Frames were collected approximately every 20 s.

---

**Figure S1.**
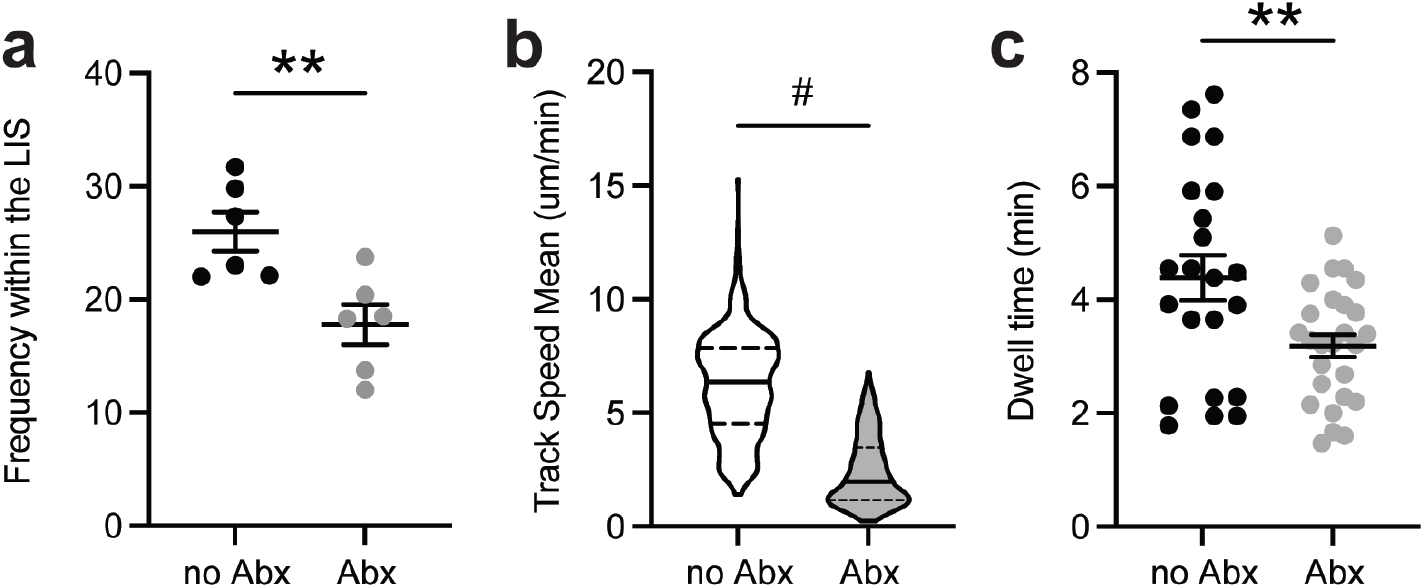
Commensal bacteria promote γδ IEL surveillance behavior. **a** Frequency of γδ IELs in the lateral intercellular space (LIS), **b** track speed mean and **c** dwell time in LIS were analyzed from intravital imaging of jejunal mucosa of untreated or antibiotic (Abx)-treated TcrdEGFP mice. Data are from two independent experiments, n=6. Each data point represents an individual mouse **a** or track **b,c.** Data represent mean (±SEM). Statistical analysis: unpaired t-test. **P<0.01, # P<0.0001.

**Figure S2.**
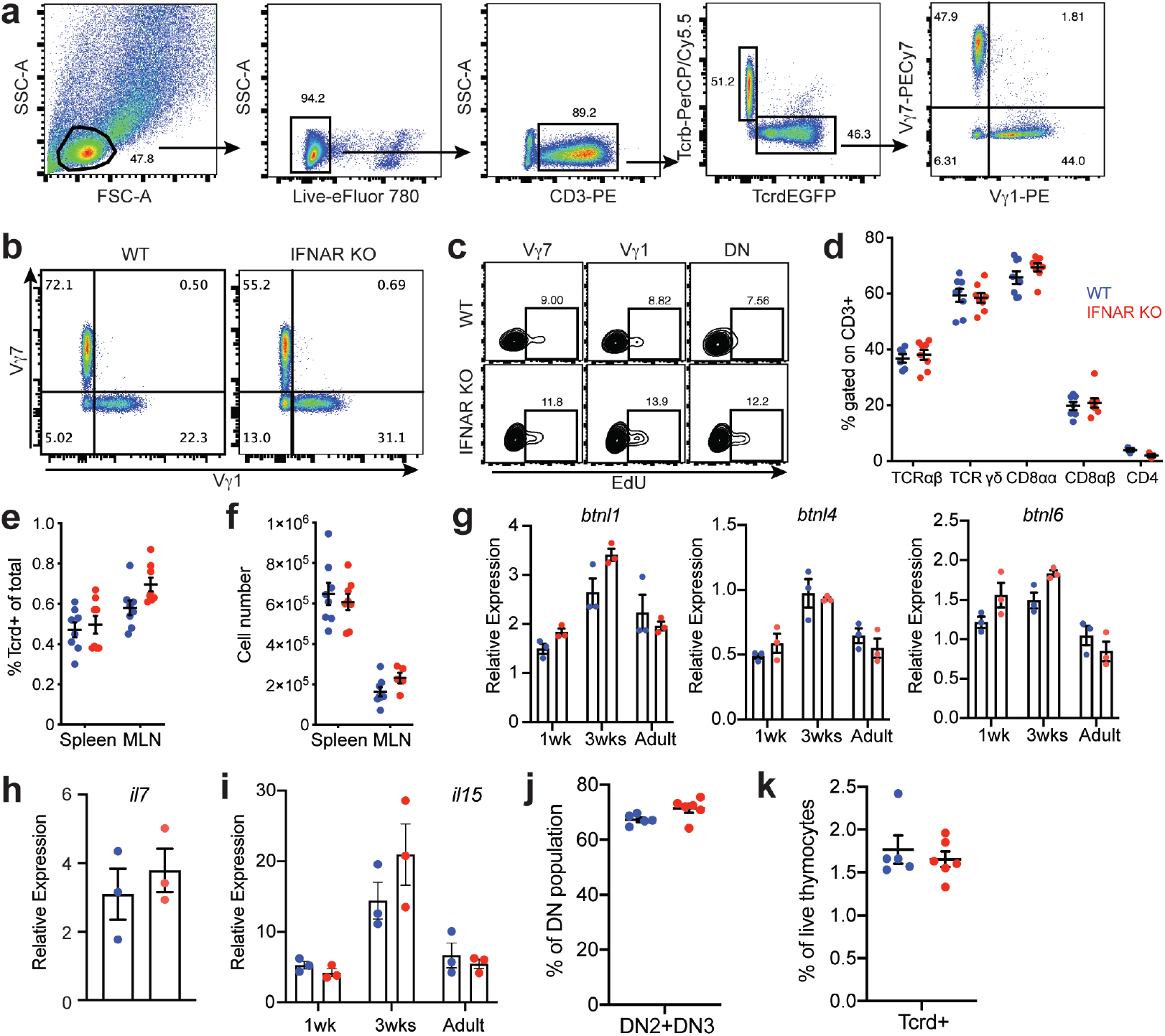
Loss of IFNAR does not alter peripheral γδ T cells, thymic γδ development or gut-intrinsic factors involved in IEL composition. **a** Gating strategy for the identification of IEL subsets. **b** Representative plot of Vγ1^+^ and Vγ7^+^ IEls and **c** EdU^+^ populations within Vγ1^+^, Vγ7^+^, and double negative (ON) IEL populations in WT-Sand IFNAR KO-S mice. **d** Phenotyping of small intestinal IEL compartment, n=6-8. **e** Frequency of Tcrd^+^ and **f** total number of γδ T cells in spleen and mesenteric lymph nodes (MLN), n=6-8. **g** Relative expression of butyrophilin-like (btnl) 1,4,6, **h** interleukin (IL)-7 or **i** IL-15 in full thickness jejunum from WT and IFNAR KO mice as measured by quantitative real-time PCR, n=3. Frequency of **j** CD25^+^ DN2 and DN3 T cell precursors and **k** Tcrd^+^ cells gated on live thymocytes in E18.5 WT-S and IFNAR KO-S mice. Data represent the mean (±SEM) from 2-3 independent experiments. Each data point represents an individual mouse. Statistical analysis: unpaired t-test.

**Figure S3.**
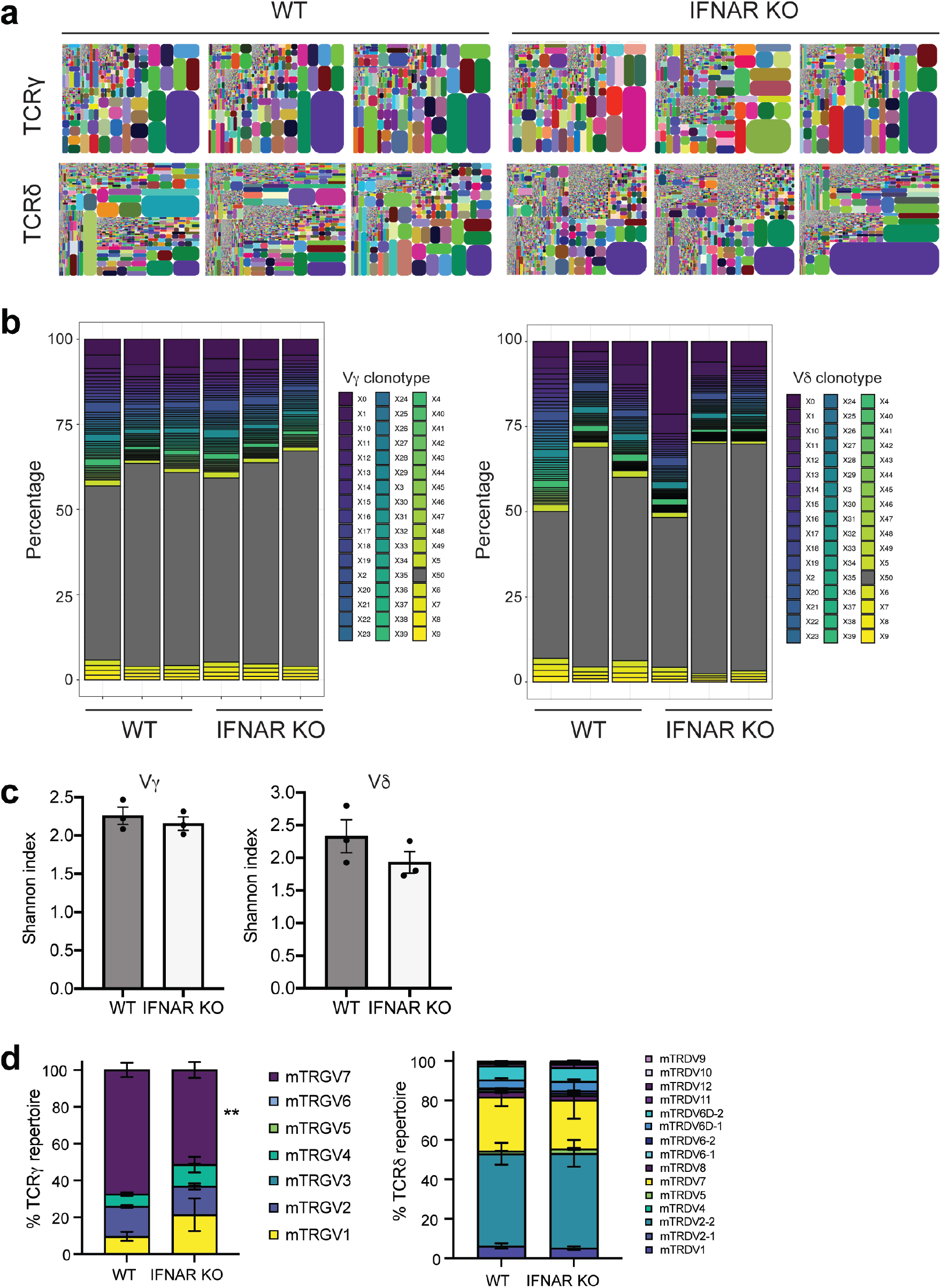
IFNAR-deficient γδ, IELs exhibit comparable TCRγδ, repertoires. **a** Representative tree maps of CDR3 clonotype usage in γδ IEls sorted from WT-S and IFNAR KO-S mice, n=3. **b** Frequency of CDR3 length usage for Vγ or Vδ subsets. **c** Shannon index of TCR diversity. **d** Percentage of TCRγ and TCRδ chain usage within the repertoire. Data represent mean (±SEM). Statistical analysis: **c:** unpaired t-test; **d:** two-way ANOVA with Sidak’s post hoc test.

**Figure S4.**
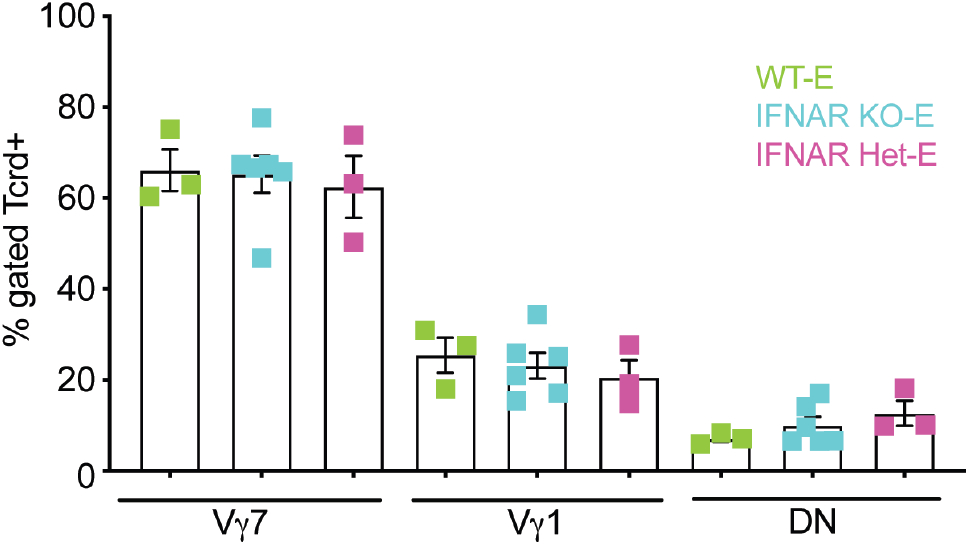
Vδ subsets are unchanged in IEls isolated from IFNAR KO-E mice. Frequency of Vγ subsets in IEL compartment from IFNAR KO-E mice. Data represent the mean (±SEM) from two independent experiments. Each data point represents an individual mouse. Statistical analysis: unpaired t-test compared to separately housed WT-S values.

**Figure S5.**
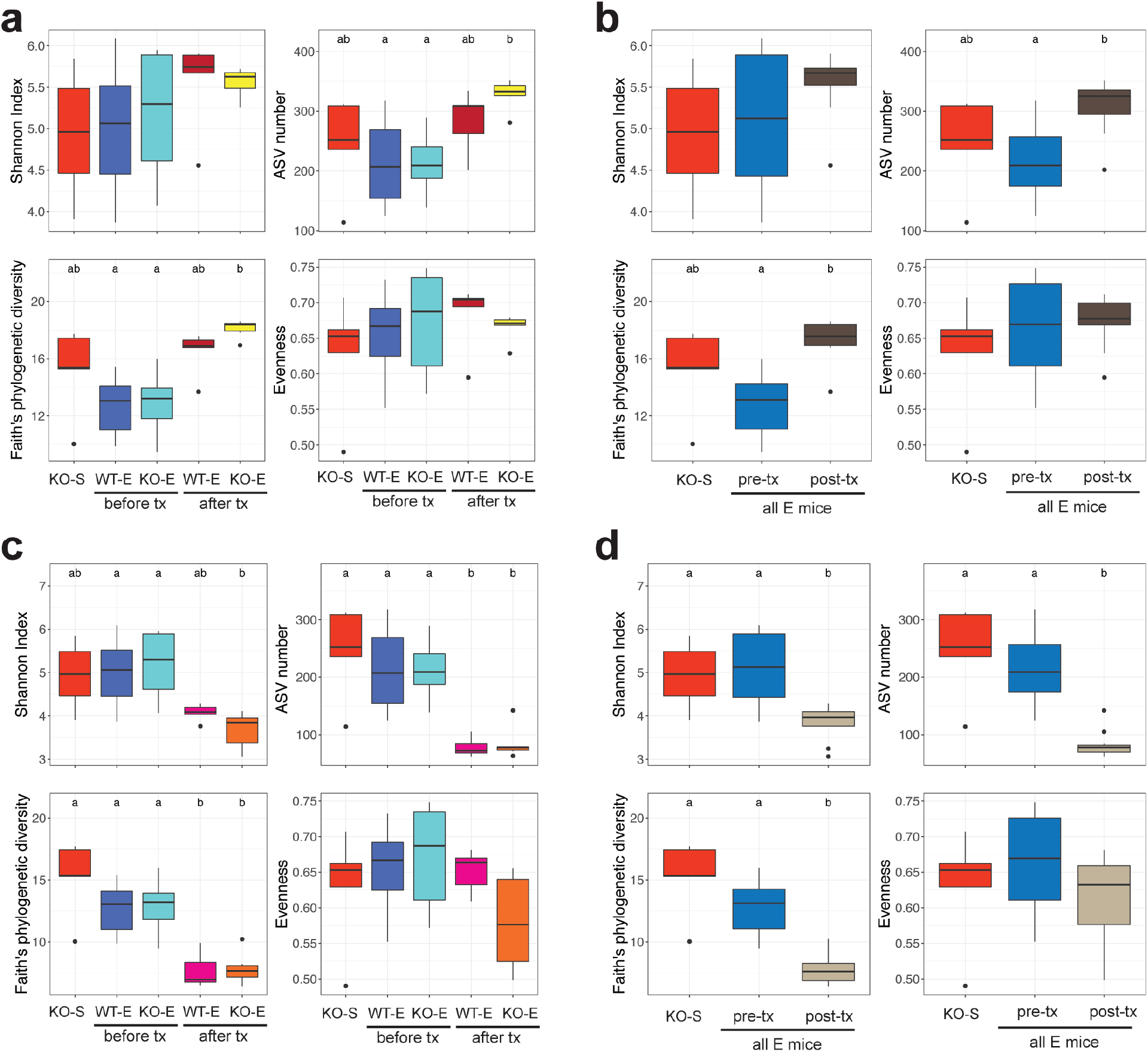
Microbial diversity is altered following horizontal transfer of IFNAR KO-S microbiota. Difference in alpha diversity of the gut microbiota between the different treatment groups. **a** Between IFNAR KO-S (donor), WT-E (pre-transfer), IFNAR KO-E (pre-transfer), WT-E (post-transfer) and IFNAR KO-E (post-transfer) **b** Between IFNAR KO-S, all E mice (pre-transfer) and all E mice (post-transfer) groups **c** Between IFNAR KO-S, WT-E (pre-transfer), IFNAR KO-E (pre-transfer), WT-E offspring (post-transfer) and IFNAR KO-E offspring (posttransfer) **d** Between IFNAR KO-S, all E mice (pre-transfer) and all E offspring mice (post-transfer) groups. Shannon Index, ASV numbers, Faith’s phylogenetic diversity and Evenness were used as the indices to reflect the alpha diversity. Statistical analysis: Kruskal-Wallis test followed by Dunn’s post hoc. Compact letters were used to show the significant differences between the groups. P<0.05 considered as significant.

**Figure S6.**
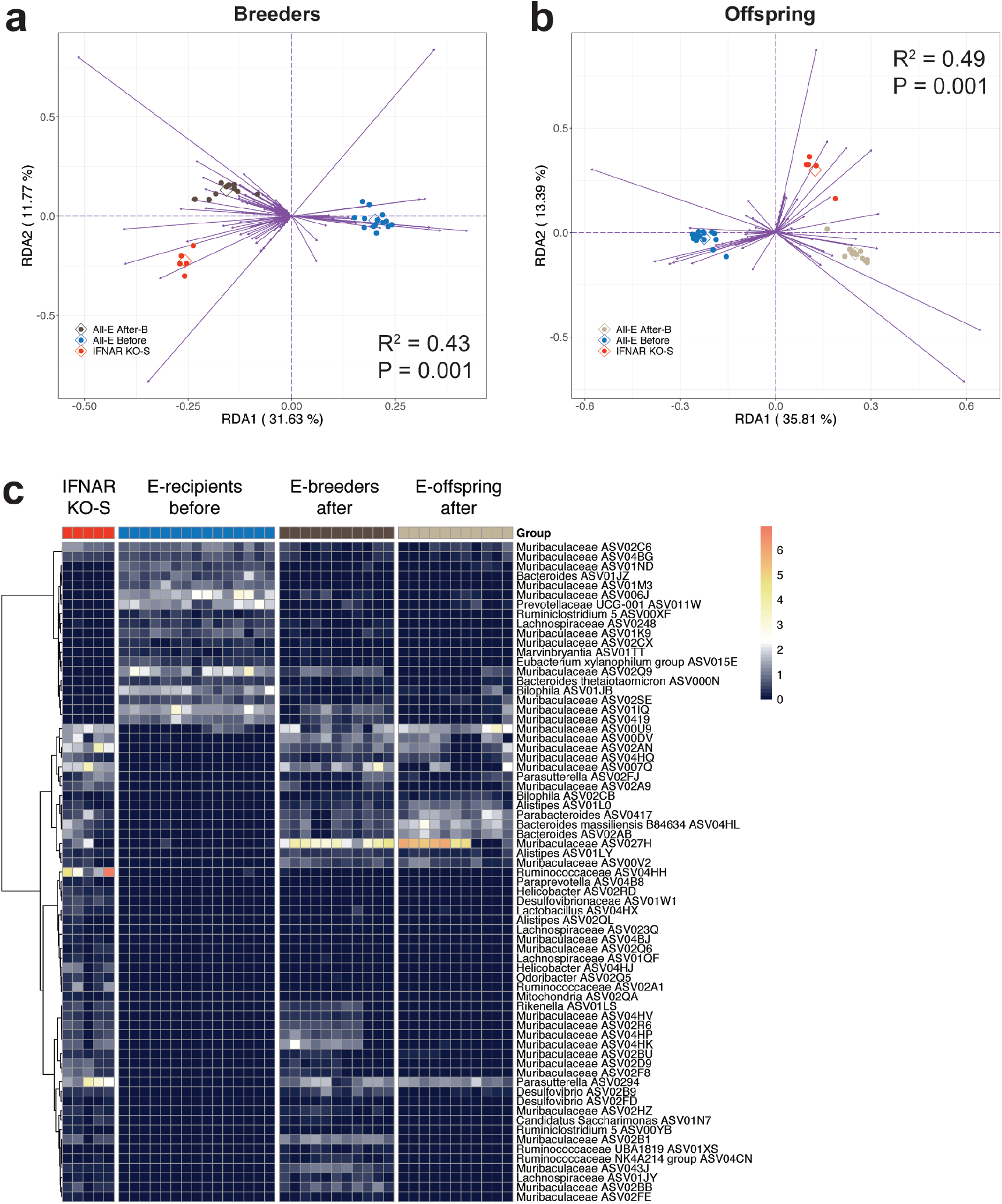
Differential microbial members are identified by redundancy analysis. Tri-plot of redundancy analysis (RDA) of the microbiota in the datasets containing **a** IFNAR KO-S, all E (pre-transfer) and all E breeders (post-transfer) groups and **b** IFNAR KO-S, all E (pre-transfer) and all E offspring (post-transfer) groups. Group information was used as the environmental variable. Hellinger transformed abundance of the (amplicon sequence variants) ASVs were used in the RDA analysis. Samples are indicated by dots. ASVs with more than 50% of the variability in their abundance explained by RDA1 and RDA2 are indicated by purple arrows. **c** The abundance heatmap of the 69 ASVs identified in both **a** and **b.** The heatmap shows the square-root transformed relative abundance of the ASVs. ASVs were clustered by ward linkage based on Spearman’s correlation coefficient.

**Figure S7.**
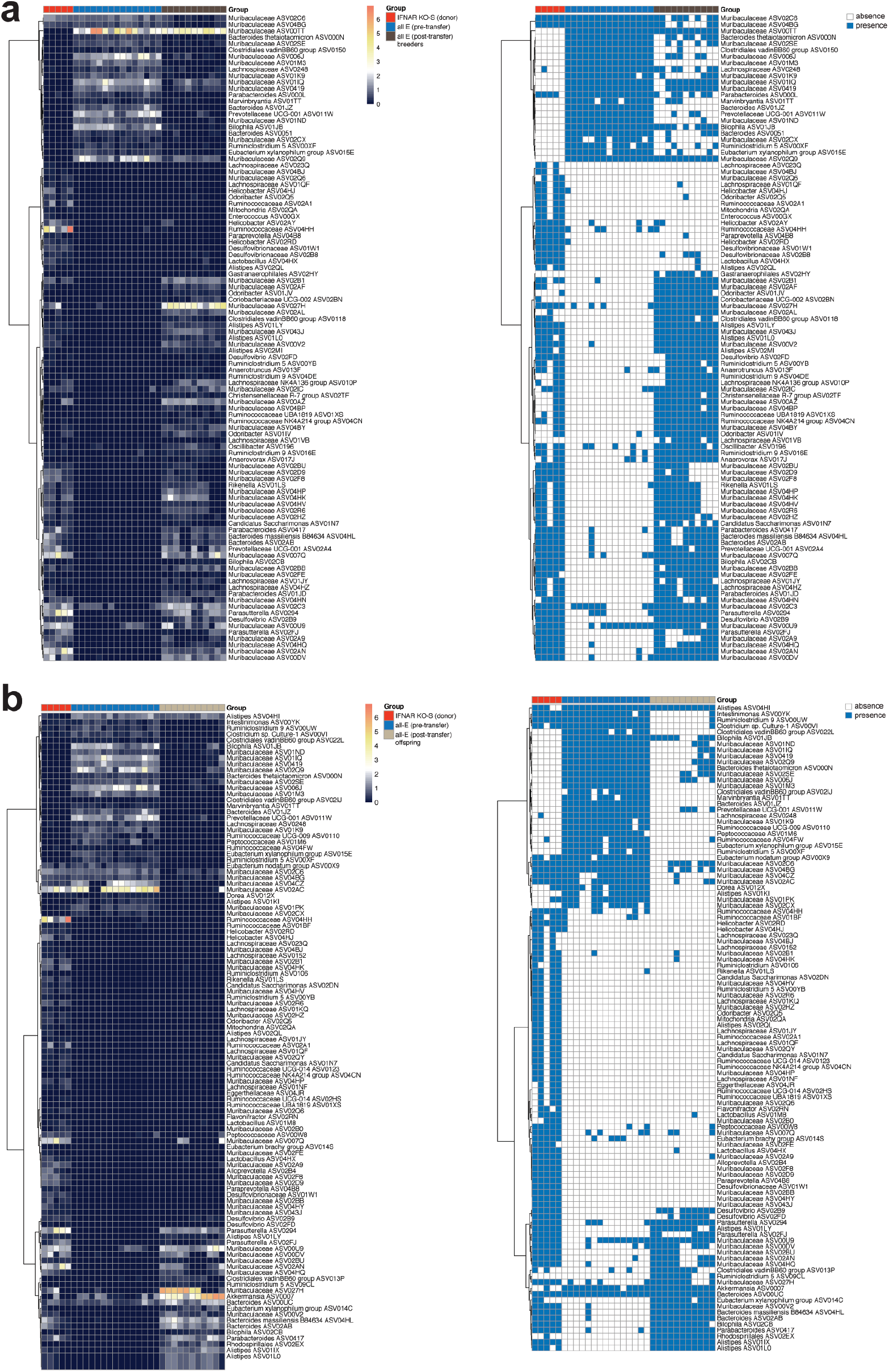
Heatmaps showing the detail abundance and occurrence information of the ASVs as identified by redundancy analysis. **a** The abundance heatmap shows the square-root transformed relative abundance **of the** 101 ASVs identified in Fig. S6a. **b** The abundance heatmap shows the square-root transformed relative abundance of the 108 ASVs identified in Fig. S6b. ASVs were clustered by ward linkage based on Spearman’s correlation coefficient. Blue and white colors in occurrence heatmap show the presence and absence of the ASVs respectively.

**Figure S8.**
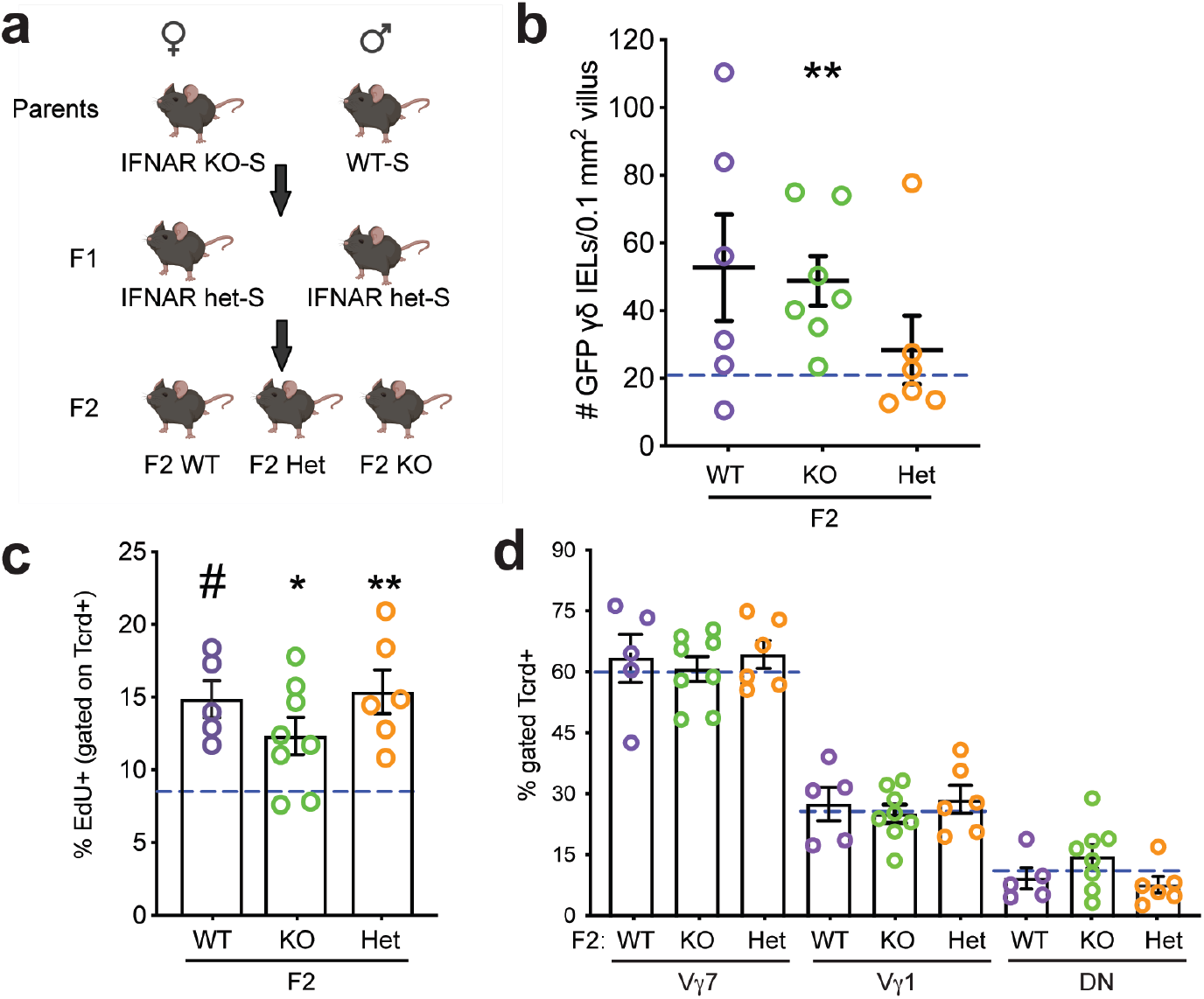
The genotype of the dam does not affect vertical transmission of the γδ IEL hyperproliferative phenotype. **a** Breeding strategy used to generate F2 WT and IFNAR KO littermates. **b** Morphometric analysis of GFP^+^ γδ T cells in the jejunum of adult F2 littermates. **c** Percentage of EdU^+^ γδ IELs and **d** proportion of Vδ IEL subsets in adult F2 littermates. Dashed lines indicate separately housed WT-S values. n=5-8 mice. Data represent mean (±SEM) from two independent experiments. Each data point represents an individual mouse. Statistical analysis: **b,c:** unpaired t-test; **d:** one-way ANOVA with Dunnett’s post hoc test; All comparisons were made to separately housed WT-E values. *P<0.05, **P<0.01, #P<0.0001.

**Figure S9.**
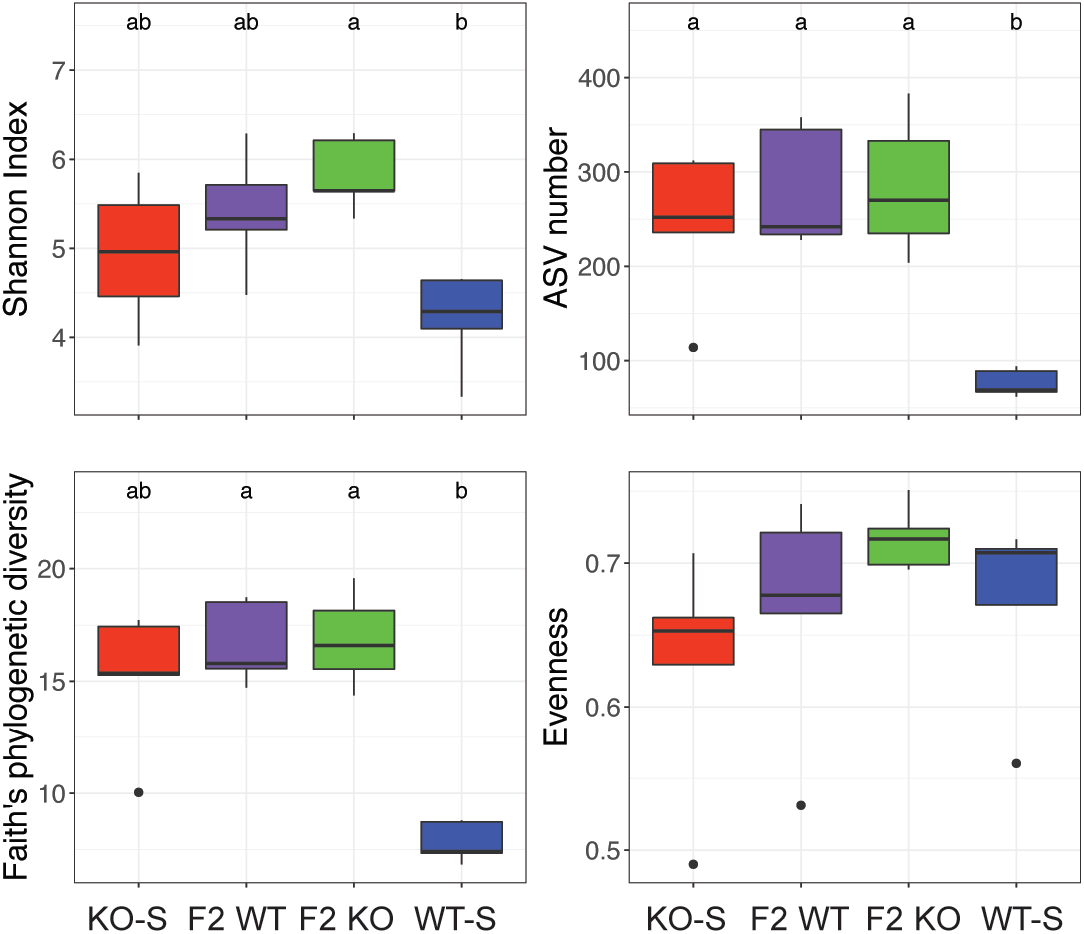
Microbial diversity is altered following vertical transfer of IFNAR KO-S microbiota. Difference in alpha diversity of the gut microbiota between the different treatment groups in the vertical transmission dataset. Shannon Index, ASV numbers, Faith’s phylogenetic diversity and Evenness were used as the indices to reflect the alpha diversity. Statistical analysis: Kruskal-Wallis test followed by Dunn’s post doc. Compact letters were used to show significance. P<0.05

## References

1. Hu MD, Jia L, Edelblum KL. Policing the intestinal epithelial barrier: Innate immune functions of intraepithelial lymphocytes. Curr Pathobiol Rep 2018; 6(1): 35–46.

2. Komano H, Fujiura Y, Kawaguchi M, Matsumoto S, Hashimoto Y, Obana S et al. Homeostatic regulation of intestinal epithelia by intraepithelial gamma delta T cells. Proc Natl Acad Sci U S A 1995; 92(13): 6147–6151.

3. Chen Y, Chou K, Fuchs E, Havran WL, Boismenu R. Protection of the intestinal mucosa by intraepithelial gamma delta T cells. Proc Natl Acad Sci U S A 2002; 99(22): 14338–14343.

4. Swamy M, Abeler-Dorner L, Chettle J, Mahlakoiv T, Goubau D, Chakravarty P et al. Intestinal intraepithelial lymphocyte activation promotes innate antiviral resistance. Nat Commun 2015; 6: 7090.

5. Ismail AS, Severson KM, Vaishnava S, Behrendt CL, Yu X, Benjamin JL et al. {gamma}{delta} intraepithelial lymphocytes are essential mediators of host-microbial homeostasis at the intestinal mucosal surface. Proc Natl Acad Sci U S A 2011; 108(21): 8743–8748.

6. Carding SR, Egan PJ. Gammadelta T cells: functional plasticity and heterogeneity. Nat Rev Immunol 2002; 2(5): 336–345.

7. Edelblum KL, Shen L, Weber CR, Marchiando AM, Clay BS, Wang Y et al. Dynamic migration of gammadelta intraepithelial lymphocytes requires occludin. Proc Natl Acad Sci U S A 2012; 109(18): 7097–7102.

8. Edelblum KL, Singh G, Odenwald MA, Lingaraju A, El Bissati K, McLeod R et al. gammadelta Intraepithelial Lymphocyte Migration Limits Transepithelial Pathogen Invasion and Systemic Disease in Mice. Gastroenterology 2015; 148(7): 1417–1426.

9. Cho H, Kelsall BL. The role of type I interferons in intestinal infection, homeostasis, and inflammation. Immunol Rev 2014; 260(1): 145–167.

10. Schaupp L, Muth S, Rogell L, Kofoed-Branzk M, Melchior F, Lienenklaus S et al. Microbiota-Induced Type I Interferons Instruct a Poised Basal State of Dendritic Cells. Cell 2020; 181(5): 1080–1096 e1019.

11. Ganal SC, Sanos SL, Kallfass C, Oberle K, Johner C, Kirschning C et al. Priming of natural killer cells by nonmucosal mononuclear phagocytes requires instructive signals from commensal microbiota. Immunity 2012; 37(1): 171–186.

12. Abt MC, Osborne LC, Monticelli LA, Doering TA, Alenghat T, Sonnenberg GF et al. Commensal bacteria calibrate the activation threshold of innate antiviral immunity. Immunity 2012; 37(1): 158–170.

13. Kernbauer E, Ding Y, Cadwell K. An enteric virus can replace the beneficial function of commensal bacteria. Nature 2014; 516(7529): 94–98.

14. Bandeira A, Mota-Santos T, Itohara S, Degermann S, Heusser C, Tonegawa S et al. Localization of gamma/delta T cells to the intestinal epithelium is independent of normal microbial colonization. J Exp Med 1990; 172(1): 239–244.

15. Suzuki H, Jeong KI, Itoh K, Doi K. Regional variations in the distributions of small intestinal intraepithelial lymphocytes in germ-free and specific pathogen-free mice. Exp Mol Pathol 2002; 72(3): 230–235.

16. Jung J, Surh CD, Lee YJ. Microbial Colonization at Early Life Promotes the Development of Diet-Induced CD8alphabeta Intraepithelial T Cells. Mol Cells 2019; 42(4): 313–320.

17. Jiang W, Wang X, Zeng B, Liu L, Tardivel A, Wei H et al. Recognition of gut microbiota by NOD2 is essential for the homeostasis of intestinal intraepithelial lymphocytes. J Exp Med 2013; 210(11): 2465–2476.

18. Liu L, Gong T, Tao W, Lin B, Li C, Zheng X et al. Commensal viruses maintain intestinal intraepithelial lymphocytes via noncanonical RIG-I signaling. Nature Immunology 2019; 20(12): 1681–1691.

19. Li Y, Innocentin S, Withers DR, Roberts NA, Gallagher AR, Grigorieva EF et al. Exogenous stimuli maintain intraepithelial lymphocytes via aryl hydrocarbon receptor activation. Cell 2011; 147(3): 629–640.

20. Cervantes-Barragan L, Chai JN, Tianero MD, Di Luccia B, Ahern PP, Merriman J et al. Lactobacillus reuteri induces gut intraepithelial CD4(+)CD8alphaalpha(+) T cells. Science 2017; 357(6353): 806–810.

21. Hoytema van Konijnenburg DP, Reis BS, Pedicord VA, Farache J, Victora GD, Mucida D. Intestinal Epithelial and Intraepithelial T Cell Crosstalk Mediates a Dynamic Response to Infection. Cell 2017; 171(4): 783–794.

22. Di Marco Barros R, Roberts NA, Dart RJ, Vantourout P, Jandke A, Nussbaumer O et al. Epithelia Use Butyrophilin-like Molecules to Shape Organ-Specific gammadelta T Cell Compartments. Cell 2016; 167(1): 203–218 e217.

23. Baccala R, Witherden D, Gonzalez-Quintial R, Dummer W, Surh CD, Havran WL et al. Gamma delta T cell homeostasis is controlled by IL-7 and IL-15 together with subset-specific factors. J Immunol 2005; 174(8): 4606–4612.

24. Ma LJ, Acero LF, Zal T, Schluns KS. Trans-presentation of IL-15 by intestinal epithelial cells drives development of CD8alphaalpha IELs. J Immunol 2009; 183(2): 1044–1054.

25. Stappenbeck TS, Virgin HW. Accounting for reciprocal host-microbiome interactions in experimental science. Nature 2016; 534(7606): 191–199.

26. Hu MD, Ethridge AD, Lipstein R, Kumar S, Wang Y, Jabri B et al. Epithelial IL-15 Is a Critical Regulator of gammadelta Intraepithelial Lymphocyte Motility within the Intestinal Mucosa. J Immunol 2018; 201(2): 747–756.

27. Knoop KA, Gustafsson JK, McDonald KG, Kulkarni DH, Coughlin PE, McCrate S et al. Microbial antigen encounter during a preweaning interval is critical for tolerance to gut bacteria. Sci Immunol 2017; 2(18).

28. Ivanov, II, Atarashi K, Manel N, Brodie EL, Shima T, Karaoz U et al. Induction of intestinal Th17 cells by segmented filamentous bacteria. Cell 2009; 139(3): 485–498.

29. Constantinides MG, Link VM, Tamoutounour S, Wong AC, Perez-Chaparro PJ, Han SJ et al. MAIT cells are imprinted by the microbiota in early life and promote tissue repair. Science 2019; 366(6464).

30. Al Nabhani Z, Dulauroy S, Marques R, Cousu C, Al Bounny S, Dejardin F et al. A Weaning Reaction to Microbiota Is Required for Resistance to Immunopathologies in the Adult. Immunity 2019; 50(5): 1276–1288 e1275.

31. Baron EJ. Bilophila wadsworthia: a unique Gram-negative anaerobic rod. Anaerobe 1997; 3(2-3): 83–86.

32. Qiu Y, Pu A, Zheng H, Liu M, Chen W, Wang W et al. TLR2-Dependent Signaling for IL-15 Production Is Essential for the Homeostasis of Intestinal Intraepithelial Lymphocytes. Mediators Inflamm 2016; 2016: 4281865.

33. Yu Q, Tang C, Xun S, Yajima T, Takeda K, Yoshikai Y. MyD88-dependent signaling for IL-15 production plays an important role in maintenance of CD8 alpha alpha TCR alpha beta and TCR gamma delta intestinal intraepithelial lymphocytes. J Immunol 2006; 176(10): 6180–6185.

34. Wilharm A, Tabib Y, Nassar M, Reinhardt A, Mizraji G, Sandrock I et al. Mutual interplay between IL-17-producing gammadeltaT cells and microbiota orchestrates oral mucosal homeostasis. Proc Natl Acad Sci U S A 2019; 116(7): 2652–2661.

35. Wilharm A, Brigas HC, Sandrock I, Ribeiro M, Amado T, Reinhardt A et al. Microbiota-dependent expansion of testicular IL-17-producing Vgamma6(+) gammadelta T cells upon puberty promotes local tissue immune surveillance. Mucosal Immunol 2021; 14(1): 242–252.

36. Jin C, Lagoudas GK, Zhao C, Bullman S, Bhutkar A, Hu B et al. Commensal Microbiota Promote Lung Cancer Development via gammadelta T Cells. Cell 2019; 176(5): 998–1013 e1016.

37. Stacy A, Andrade-Oliveira V, McCulloch JA, Hild B, Oh JH, Perez-Chaparro PJ et al. Infection trains the host for microbiota-enhanced resistance to pathogens. Cell 2021; 184(3): 615–627 e617.

38. Yeung F, Chen YH, Lin JD, Leung JM, McCauley C, Devlin JC et al. Altered Immunity of Laboratory Mice in the Natural Environment Is Associated with Fungal Colonization. Cell Host Microbe 2020; 27(5): 809–822 e806.

39. Rosshart SP, Vassallo BG, Angeletti D, Hutchinson DS, Morgan AP, Takeda K et al. Wild Mouse Gut Microbiota Promotes Host Fitness and Improves Disease Resistance. Cell 2017; 171(5): 1015–1028 e1013.

40. Rauch I, Hainzl E, Rosebrock F, Heider S, Schwab C, Berry D et al. Type I interferons have opposing effects during the emergence and recovery phases of colitis. Eur J Immunol 2014; 44(9): 2749–2760.

41. Tschurtschenthaler M, Wang J, Fricke C, Fritz TM, Niederreiter L, Adolph TE et al. Type I interferon signalling in the intestinal epithelium affects Paneth cells, microbial ecology and epithelial regeneration. Gut 2014; 63(12): 1921–1931.

42. Hu MD, Edelblum KL. Sentinels at the frontline: the role of intraepithelial lymphocytes in inflammatory bowel disease. Curr Pharmacol Rep 2017; 3(6): 321–334.

43. Prinz I, Sansoni A, Kissenpfennig A, Ardouin L, Malissen M, Malissen B. Visualization of the earliest steps of gammadelta T cell development in the adult thymus. Nat Immunol 2006; 7(9): 995–1003.

44. Jia L, Edelblum KL. Intravital Imaging of Intraepithelial Lymphocytes in Murine Small Intestine. J Vis Exp 2019; (148).

45. Costea PI, Zeller G, Sunagawa S, Pelletier E, Alberti A, Levenez F et al. Towards standards for human fecal sample processing in metagenomic studies. Nature Biotechnology 2017; 35(11): 1069–1076.

46. Parada AE, Needham DM, Fuhrman JA. Every base matters: assessing small subunit rRNA primers for marine microbiomes with mock communities, time series and global field samples. Environmental Microbiology 2016; 18(5): 1403–1414.

47. Apprill A, McNally S, Parsons R, Weber L. Minor revision to V4 region SSU rRNA 806R gene primer greatly increases detection of SAR11 bacterioplankton. Aquatic Microbial Ecology 2015; 75(2): 129–137.

48. Wang J, Zhang Q, Wu G, Zhang C, Zhang M, Zhao L. Minimizing spurious features in 16S rRNA gene amplicon sequencing: PeerJ Preprints; 2018. Report no. 2167-9843.

49. Quast C, Pruesse E, Yilmaz P, Gerken J, Schweer T, Yarza P et al. The SILVA ribosomal RNA gene database project: improved data processing and web-based tools. Nucleic acids research 2012; 41(D1): D590–D596.

